# Light-dependent THRUMIN1 phosphorylation regulates its association with actin filaments and 14-3-3 proteins

**DOI:** 10.1101/2021.07.09.451853

**Authors:** Matthew E. Dwyer, Roger P. Hangarter

**Affiliations:** Department of Biology, Indiana University, Bloomington, IN 47405, USA

## Abstract

Light-dependent chloroplast movements in leaf cells contribute to the optimization of photosynthesis. Low light conditions induce chloroplast accumulation along periclinal cell surfaces, providing greater access to the available light, whereas high light induces movement of chloroplasts to anticlinal cell surfaces providing photodamage protection and allowing more light to reach underlying cell layers. The THRUMIN1 protein is required for normal chloroplast movements in *Arabidopsis thaliana* and has been shown to localize at the plasma membrane and to undergo rapid light-dependent interactions with actin filaments through the N-terminal intrinsically disordered region. A predicted WASP-Homology 2 (WH2) domain was found in the intrinsically disordered region but mutations in this domain did not disrupt localization of THRUMIN1:YFP to actin filaments. A series of other protein truncations and site-directed mutations of known and putative phosphorylation sites indicated that a phosphomimetic mutation (serine to aspartic acid) at position 170 disrupted localization of THRUMIN1 with actin filaments. However, the phosphomimetic mutant rescued the *thrumin1-2* mutant phenotype for chloroplast movement and raises questions about the role of THRUMIN1’s interaction with actin. Mutation of serine 146 to aspartic acid also resulted in cytoplasmic localization of THRUMIN1:YFP in *Nicotiana benthamiana*. Mutations to a group of putative zinc-binding cysteine clusters implicates the C-terminus of THRUMIN1 in chloroplast movement. Phosphorylation-dependent association of THRUMIN1 with 14-3-3 KAPPA and OMEGA were also identified. Together, these studies provide new insights into the mechanistic role of THRUMIN1 in light-dependent chloroplast movements.

**One Sentence Summary:** Site-directed mutagenesis of THRUMIN1 revealed critical sites involved in blue light-dependent localization of THRUMIN1 to actin filaments, 14-3-3 proteins, and its regulation of chloroplast movement.

## Introduction

Plants have evolved several light-sensing mechanisms that protect chloroplasts from photodamage and for photosynthetic optimization (Li et al., 2009). Light-dependent chloroplast movements are a cellular level response to dynamic changes in light levels in their environment. In low light conditions, leaf mesophyll cells initiate what is known as the accumulation response in which the cells position their chloroplasts along the top and bottom periclinal cell faces (Zurzycki, 1955). When exposed to strong light, the cells initiate an avoidance response in which they position their chloroplasts along the anticlinal sides of the cells (Kasahara et al., 2002). In angiosperms, this mechanism is controlled by the Phototropin1 (Phot1) and Phototropin2 (Phot2) blue light photoreceptors (Jarillo et al., 2001; Kagawa et al., 2001; Sakai et al., 2001). The two photoreceptors are functionally redundant but Phot1 primarily induces the accumulation response while Phot2 induces the avoidance response (Sakai et al., 2001; Luesse et al., 2010). In addition to the Phototropins, phytochromes (red/far red photoreceptors) have been shown to play a role in modulating the response to blue light-induced chloroplast movement through inhibition of Phototropin-related pathways and aiding in the transition from low to high light responses (DeBlasio et al., 2003; Luesse et al., 2010). The importance of chloroplast movements has been demonstrated by observations of significantly more photodamage in *Arabidopsis thaliana* plants that have defective Phototropins and/or mutated forms of other proteins needed for normal chloroplast positioning, when compared to wild-type plants (Kasahara et al., 2002; Davis and Hangarter, 2012). In addition to reducing photodamage, the avoidance response in palisade mesophyll cells allows more photons to reach underlying spongy mesophyll cells and enhances whole-leaf photosynthesis (Davis and Hangarter, 2012).

Angiosperms require the actin cytoskeleton for normal chloroplast movements (Malec et al., 1996; Kandasamy and Meagher, 1999; Kadota et al., 2009). No genetic evidence for motor proteins has been seen after extensive mutant screens so research has focused on identifying how actin dynamics may drive chloroplast movement (Avisar et al., 2008; Suetsugu et al., 2010; Wada, 2013). Using the actin-binding probe, GFP:mTalin, actin filaments have been shown to associate with the chloroplast outer envelope in a light-dependent way (Kadota et al., 2009). Upon irradiation with high intensity blue light, the GFP:mTalin was found to disappear from the chloroplast-associated actin filaments (cp-actin) suggesting disassociation of cp-actin on the time scale of minutes. Upon removal of the blue light stimulus, GFP:mTalin-labeled cp-actin was found to reappear (Kadota et al., 2009). Moreover, when only a portion of a cell was exposed to blue light, the chloroplasts in the irradiated area lost their GFP:mTalin-labeled cp-actin but as the chloroplasts were leaving the lit region of a cell, GFP:mTalin-labeled cp-actin was seen to reappear along the leading edge of movement. The Phot2-dependent dynamics of cp-actin are thought to be critical to the mechanism that drives the chloroplast movements (Kong et al., 2013).

THRUMIN1 is a light-dependent, plasma membrane-localized F-actin bundling protein and loss of THRUMIN1 was found to result in reduced chloroplast motility suggesting a role for THRUMIN1 bundling of F-actin at the plasma membrane (Whippo et al., 2011). THRUMIN1 requires myristoylation and/or the palmitoylation at the N-terminus for proper localization to filamentous actin and for chloroplast movements (Whippo et al., 2011). The intrinsically disordered region (IDR) of THRUMIN1 was previously shown to be the region that confers F-actin binding (Whippo et al., 2011) but the IDR domain alone failed to rescue chloroplast movements in the *thrumin1-2* mutant. Expression of just the C-terminal region also failed to rescue the mutant phenotype and showed diffuse, non-filamentous localization (Whippo et al., 2011). THRUMIN1 is conserved for its glutaredoxin-like domain but the IDR and the C-terminal cysteine-rich region appear to be the critical regions for the light-induced bundling activity of THRUMIN1 (Whippo et al., 2011).

The requirement of F-actin for chloroplast movement is undisputed but the signaling cascade and motility mechanism that regulates light-dependent chloroplast movement remain unknown. Given that Phot1 and Phot2 are blue light receptor kinases (Liscum and Briggs, 1995; Briggs and Huala, 1999), phosphorylation is likely to play a role in regulating chloroplast movement. Phosphoproteomic analyses of plasma membrane associated proteins showed that THRUMIN1 was among the proteins found to undergo light-dependent phosphorylation (Boex-Fontvieille et al., 2014). Specifically, serine 113 or 115 and serine 164 were found to be more phosphorylated in the dark and mutations of those residues were reported to have altered light-dependent chloroplast movement (Boex-Fontvieille et al., 2014).

The schematic view of THRUMIN1 shown in Fig. 1 shows its major domains and predicted sites of phosphorylation, 14-3-3 protein recognition motifs, a putative WH2 domain, and the location of several putative zinc-binding cysteine residues. Here we report the results of a number of new site-directed mutants and protein truncations of THRUMIN1 that focused on the potential roles of phosphorylation, protein-protein binding, and the putative WH2 domain in THRUMIN1’s ability to localize to F-actin and modulate light-dependent chloroplast movements. We found specific phosphorylation-dependent interactions between THRUMIN1 and 14-3-3 KAPPA and OMEGA. The predicted WH2 domain did not appear to be the site of actin binding but mutation of two putative phosphorylation sites near the WH2 domain disrupted THRUMIN1-actin filament localization without interfering with chloroplast movement dynamics suggesting that THRUMIN1’s interaction with cp-actin may not be directly associated with the motility mechanism as previously thought. These and other new results reported in this paper lead us to propose that THRUMIN1 may function in anchoring chloroplasts to the plasma membrane rather than being part of the motive force for chloroplast movements. Overall, our studies provide new insights into the role of THRUMIN1 phosphorylation and protein associations involved in the molecular mechanism of chloroplast movement.

**Figure 1.**
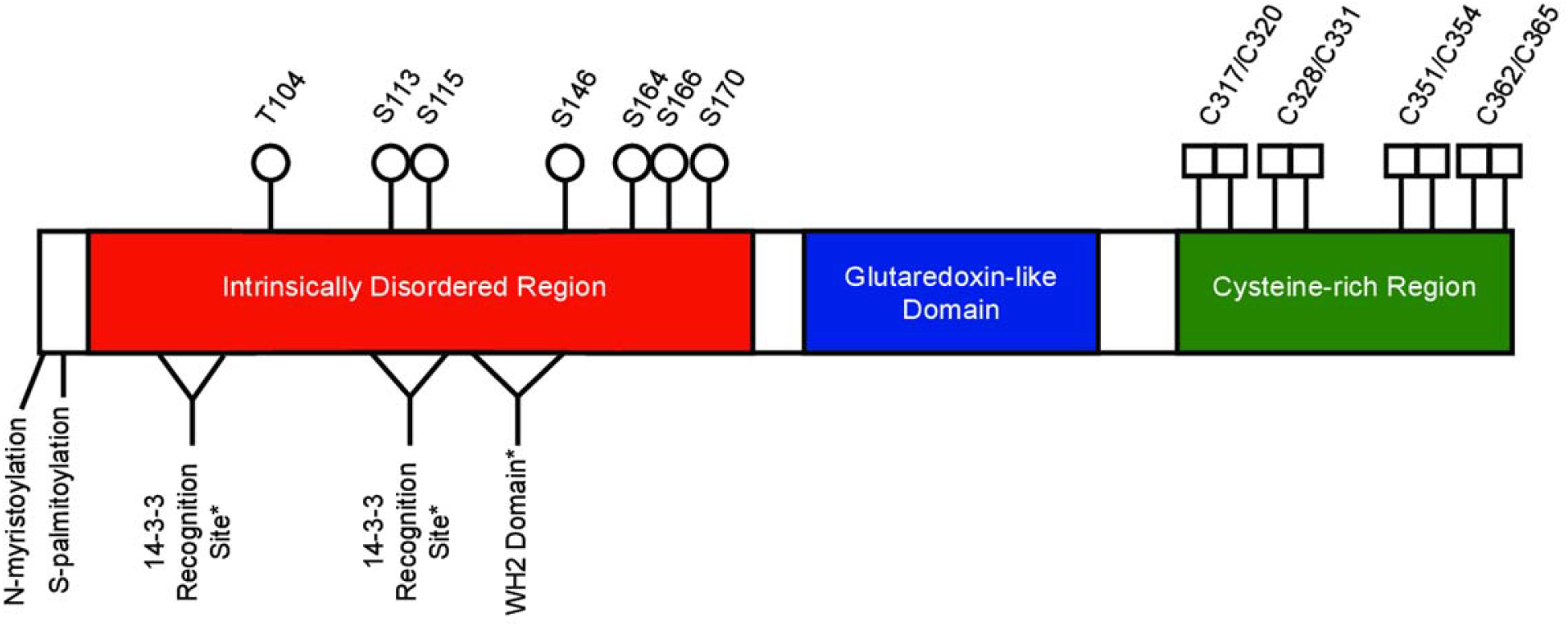
Schematic of THRUMIN1 showing known and predicted sites of interest. Previously known sites in the intrinsically disordered region (red) include the N-myristoylation and S-palmitoylation (plasma membrane tethering) and phosphorylation sites S113, S115, S164, and S166. S113, S115, S164, and S166 were previously reported as phosphorylation sits by (Boex-Fontvieille et al., 2014). Phosphorylation sites, T104 and S146, were identified in this study by mass spectrometry. Serine 170 is a putative phosphorylation site that we found in this study to be important for THRUMIN1 function. Two putative 14-3-3 recognition sites (amino acids 50-54 and 111-117) and a putative WASP Homology-2 (WH2) actin-binding domain (amino acids 127-144) were predicted by the Eukaryotic Linear Motif program. Putative zinc-binding cysteine residues are shown in the cysteine-rich region (green). The glutaredoxin-like domain (blue), has no known function in chloroplast movement (Whippo et al., 2011).

## Results

### THRUMIN1 dynamically associates with chloroplast-associated actin filaments

THRUMIN1 is a plasma-membrane associated protein with actin-bundling activity and is required for normal light-dependent chloroplast movements in *Arabidopsis thaliana* (Whippo et al., 2011). Our time-lapse studies of the dynamics of THRUMIN1:YFP localization in palisade mesophyll cells showed that it associates with actin filaments at the periphery of the chloroplast envelope in a light-dependent manner (Fig. 2) similar to what has been observed with the actin-binding GFP:mTalin probe (Kadota et al., 2009; Kong et al., 2013). After dark acclimation, THRUMIN1 was localized around the periphery of the chloroplast outer envelope but upon light stimulation of a portion of a cell, THRUMIN1 rapidly dissociated from the chloroplast periphery and then reassociated at the leading edge of the chloroplasts when they began to move towards areas of the cell away from the blue light (Fig. 2, Supplemental Fig. S1A). The difference between the THRUMIN1 fluorescence on the leading edge versus the lagging edge of the chloroplast was significant (p=0.0291) at peak movement of the chloroplast (Supplemental Fig. S1B). The dynamics of blue light-induced THRUMIN1 relocalization can be seen in the Supplemental Movies S1 and S2.

**Figure 2.**
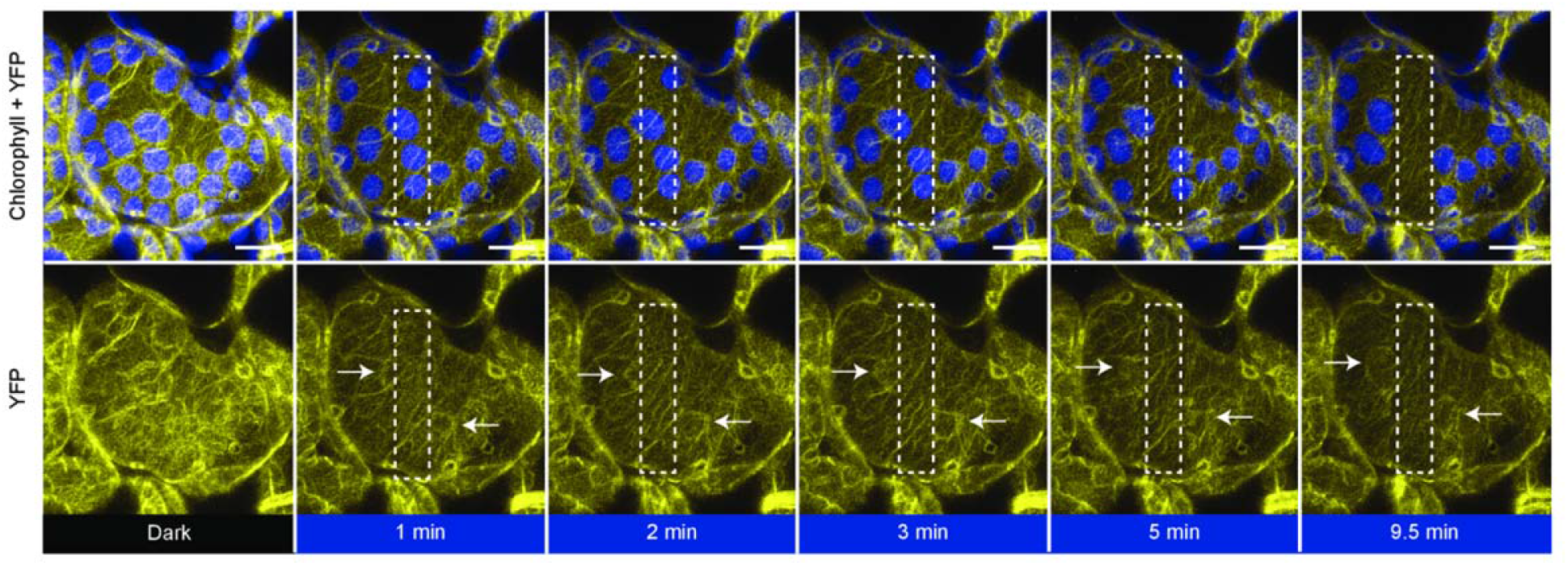
THRUMIN1:YFP dynamically localizes to cp-actin filaments in a light-dependent manner. In a dark state (YFP excitation, 514nm), 35S:THRUMIN1:YFP expressed in *thrumin1-2 A. thaliana* cells displayed a uniform, basket-like organization around the chloroplast periphery. Within a minute of blue light stimulation (white rectangle), THRUMIN1:YFP disappeared and reorganized towards the leading edge of movement as indicated by the white arrows in the YFP channel. After 10 minutes of blue light stimulation, the chloroplasts inside the white rectangle have exited the region and the THRUMIN1:YFP filaments started to become more peripheral around the chloroplasts. Chlorophyll autofluorescence is false-colored blue and the YFP channel is separated. Representative time-lapse images are shown for the dark treatment (514nm for YFP excitation) and blue light stimulation intervals (470nm and 514nm). The scale bar indicates a 5 µm distance.

### The chloroplast outer envelope displays amoeboid-like movements

Most research on chloroplast movements has relied on chlorophyll autofluorescence for following their movements and associations with other proteins. In a study by Kong et al (2013) they used the chloroplast membrane marker OEP7:YFP and showed a correlation between chloroplast movements and movement of chloroplast envelope membranes away from the thylakoid membrane (Kong et al., 2013). In this study, we made use of the greater contrast available between chlorophyll autofluorescence and the stroma-localized probe tpFNR:YFP to observe chloroplast membrane dynamics and found that the chloroplast envelope exhibited highly dynamic, pseudopod-like protrusions that appear to be associated with chloroplast movements (Fig 3A and Supplemental Movies S3 and S4). The membrane protrusions became more dynamic in response to localized blue-light exposure and, consistent with Kong et al. (2013), the protrusions quickly became more pronounced near the leading edge of moving chloroplasts and corresponded to where THRUMIN1 was also found to localize after light stimulation (Fig. 2, Fig. 3A,C, and Supplemental Movies S1, S2, S3, and S4). Upon removal of the blue light stimulus, the protrusions became more distributed around the chloroplast periphery (Fig 3A and Supplemental Movies S3 and S4). In *thrumin1-2* mutant plants expressing the tpFNR:YFP transgene, membrane protrusions were not observed and the movement of chloroplasts was more erratic than with wild type THRUMIN1 (Fig. 3B,C, Supplemental Movies S5 and S6). Time-lapse movies of the pseudopod-like chloroplast membrane protrusions in wild-type plants evoke an amoeboid-like crawling behavior. However, unlike the internal forces of actin dynamics that drive pseudopod extension in amoeba, chloroplast actin is external and suggests that THRUMIN1 has a role in anchoring cp-actin filaments to the plasma membrane.

**Figure 3.**
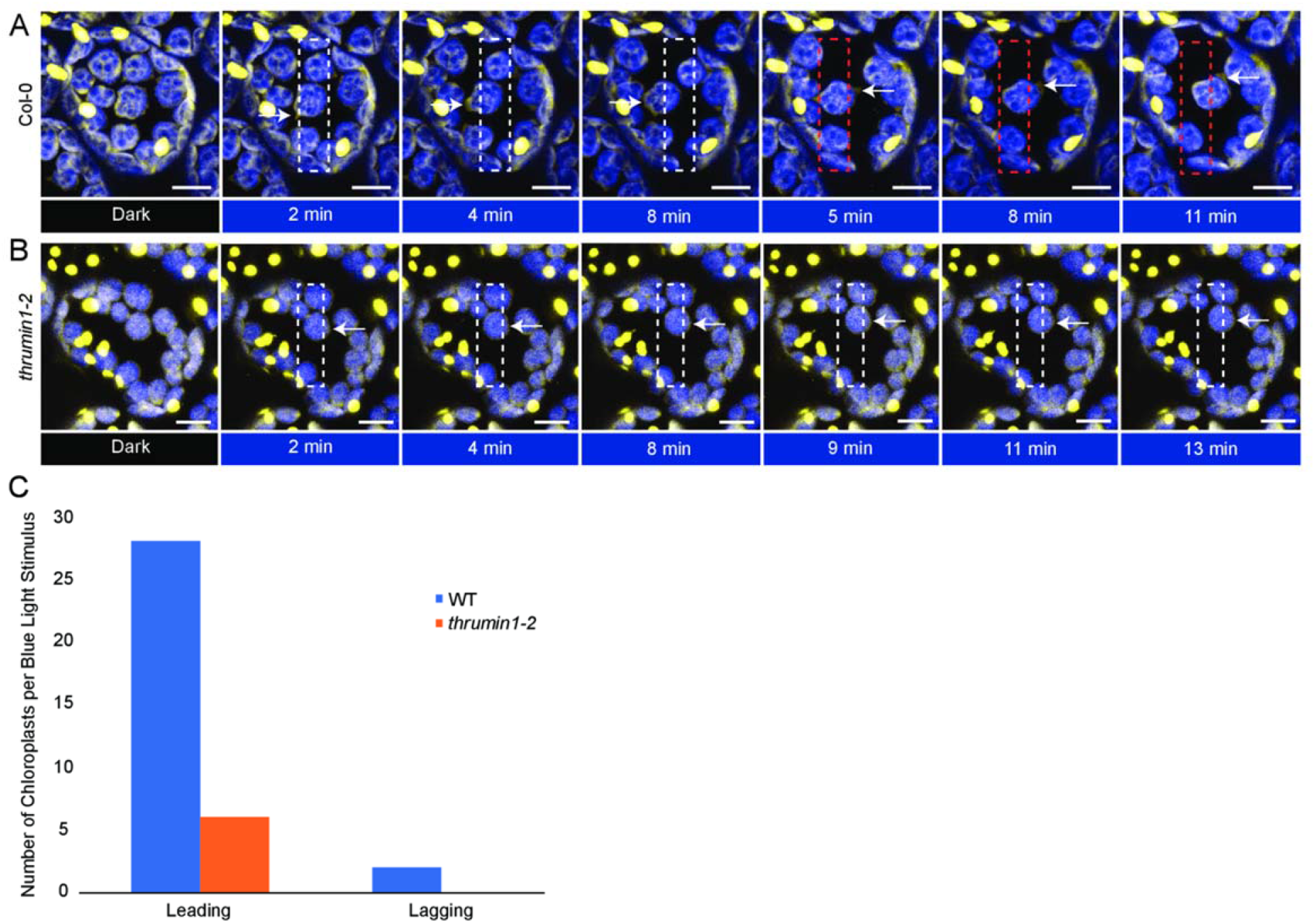
Chloroplast membranes display directional amoeboid-like movements in response to blue light. (A) Col-0 wild type plants expressing the stroma-localized marker tpFNR:YFP were imaged in a dark state (YFP excitation, 514nm) and a blue light-stimulated region of interest (white rectangle, 470nm). As per normal chloroplast movement kinetics, within a minute of blue light stimulation the chloroplasts began to leave the region of interest with a membrane protrusion in the direction of movement (white arrows). When the region of blue light stimulation is altered to force a new direction of chloroplast movement (red rectangle), a new membrane protrusion in the direction of movement formed. (B) *thrumin1-2* mutant plants expressing the tpFNR:YFP marker did not exhibit robust membrane protrusions (white arrows) in response to a blue light stimulus (white rectangle). The chloroplasts moved in slow but sporadic patterns with no discernable membrane activity after 13 minutes of a blue light stimulus. Chlorophyll autofluorescence is false-colored blue and the YFP channel is false-colored yellow. Representative time-lapse images are shown for the dark treatment (514nm for YFP excitation) and blue light stimulation intervals (470nm and 514nm). The scale bar indicates a 5 µm distance. (C) THRUMIN1 association with the development of chloroplast membrane protrusions on the leading edge of chloroplasts in response to high blue light microbeam irradiation. Chloroplast membrane protrusion events were counted along the leading and lagging edges in wild type and *thrumin1-2* mutant backgrounds. The histograms show the total leading/lagging edge protrusion events for 31 chloroplasts from 8 different cells for wild type and 30 chloroplasts from 9 different cells for the *thrumin1-2* mutant.

### Putative WASP Homology-2 domain encoded in THRUMIN1 does not confer localization of THRUMIN1:YFP to actin filaments

The IDR region of THRUMIN1 was previously shown to be required for its actin binding/bundling activity (Whippo et al., 2011). A predicted WASP Homology-2 (WH2) domain was identified within the IDR using The Eukaryotic Linear Motif Resource (Gouw et al., 2018) (Fig. 1). Because WH2 domains in many actin monomer- and polymer-binding proteins have been found to be important in regulating their interaction with actin (Paunola et al., 2002; Hertzog et al., 2004; Loomis et al., 2006), the predicted WH2 domain in THRUMIN1 seemed like a good candidate for THRUMIN1’s F-actin binding activity. However, when the putative WH2 domain was deleted (THRUMIN1^ΔWH2^:YFP) the protein still localized with F-actin along the chloroplast periphery when transiently expressed in *N. benthamiana* (Fig. 4B). In addition, THRUMIN1:YFP with point mutations in the conserved L141 and K142 residues of the putative WH2 domain also showed normal F-actin localization along the chloroplast periphery (Fig. 4C). Moreover, expression of the THRUMIN1^ΔWH2^:YFP transgene in *A. thaliana thrumin1-2* mutant plants fully rescued the mutant phenotype (Fig. 4D). Thus, the predicted WH2 domain does not appear to be directly involved in THRUMIN1’s interaction with F-actin nor for its function in regulating chloroplast movement.

**Figure 4.**
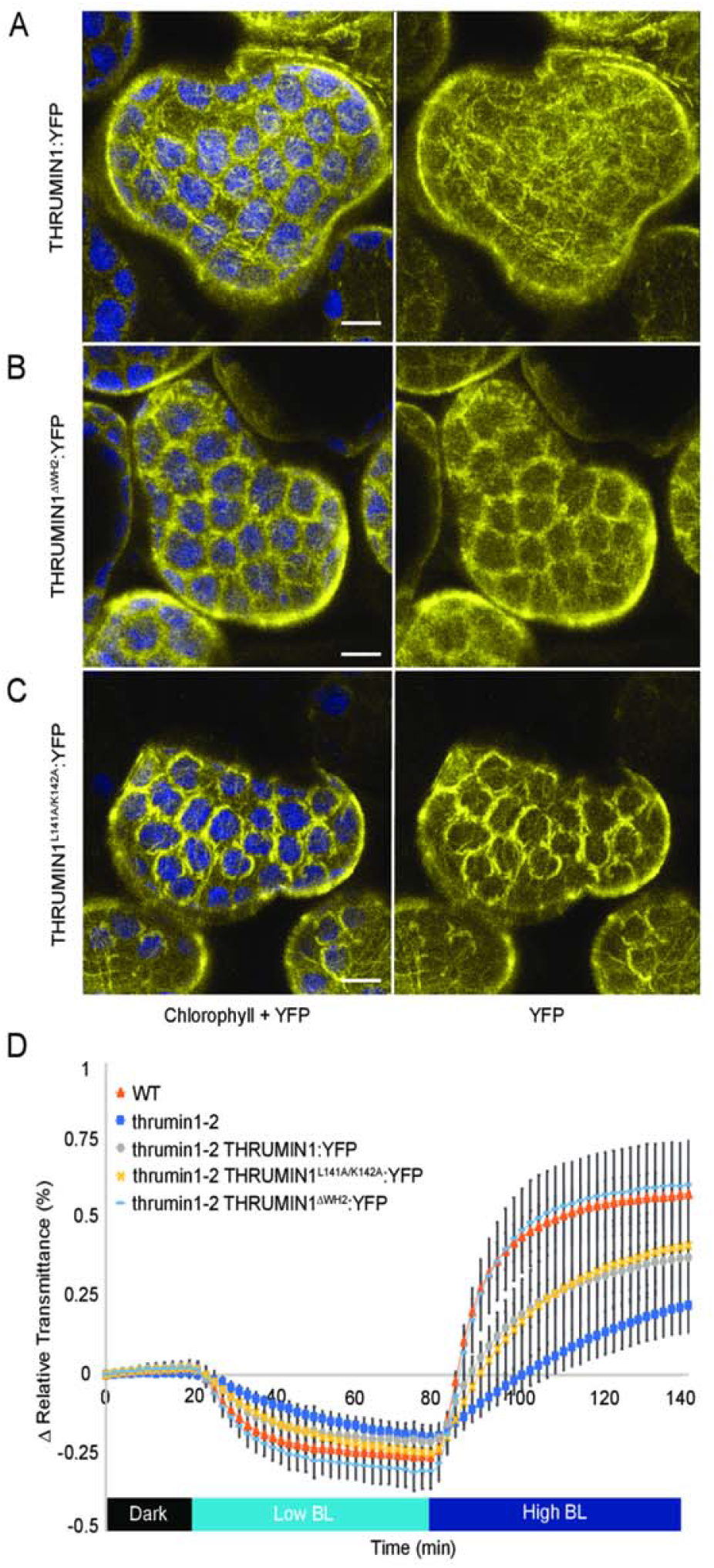
Effect of deletion of the putative WH2 domain on the localization of THRUMIN1. (A) Transient expression of wild-type 35S:THRUMIN1:YFP in *N. benthamiana* leaves showed filamentous localization around the chloroplast periphery. (B) Deletion of the WH2 domain from THRUMIN1 and (C) mutagenesis of two conserved amino acid residues (L141 and K142) in the WH2 domain did not disrupt the filamentous localization compared to wild type. Chlorophyll autofluorescence is false-colored blue and the YFP channel is separated. All of the THRUMIN1 constructs were under control of 35S promoter. The scale bar indicates a 5 µm distance. (D) Light transmittance assay of chloroplast movements in wildtype (WT) Col-0, *thrumin1-2* mutant, *thrumin1-2* 35S:THRUMIN1:YFP, *thrumin1-2* 35S:THRUMIN1^ΔWH2^:YFP, and *thrumin1-2* 35S:THRUMIN1^L141A/K142A^:YFP rescue genotypes. Deletion of the WH2 domain conferred wild type chloroplast movements. After establishment of the baseline level of leaf light transmittance after dark acclimation, chloroplast movement was induced by treatment with low blue light (∼2 µmol m^−2^s^−1^) followed by high blue light intensity (∼100 µmol m^−2^s^−1^). Standard deviation error bars represent the variance in transmittance values for 8-12 individual plants per genotype.

### Serine 170 is required for light-dependent F-actin localization of THRUMIN1:YFP

Because the deletion of the predicted WH2 domain failed to disrupt the ability of THRUMIN1 to interact with actin, a series of other internal protein truncations within the IDR (amino acids #1-201) of THRUMIN1:YFP were created and assayed for their F-actin localization when transiently expressed in *N. benthamiana*. Confocal microscopy revealed that the chloroplast-associated filamentous localization of THRUMIN1:YFP was disrupted in THRUMIN1^Δ135-201^:YFP, THRUMIN1^Δ152-201^:YFP, and THRUMIN1^Δ169-201^:YFP deletion mutants. However, the deletion mutant THRUMIN1^Δ186-201^:YFP showed normal filamentous localization (Fig. 5A). Those results indicated that localization of THRUMIN1 to cp-actin filaments resided between amino acids 169 through 186 and suggested that serine 170 may be a phosphorylation site. When a serine 170 to aspartic acid mutant form of THRUMIN1 was transiently expressed in *N. benthamiana* leaf cells, it failed to localize to filaments and conferred a more diffuse localization phenotype with or without whole-field blue light (Fig. 5B, Supplemental Movie S7). Diffuse localization of the THRUMIN1^S170D^:YFP mutant transgene was also observed with or without whole-field blue light when stably expressed in *A. thaliana thrumin1-2* mutant plants (Supplemental Movie S8). However, THRUMIN1^S170D^:YFP rescued the defective chloroplast movement phenotype in the *thrumin1-2* mutant plants (Fig. 5C, Supplemental Movie S8). When a non-phosphorylatable THRUMIN1^S170A^:YFP mutant was expressed in *N. benthamiana* and in the *A. thaliana thrumin1-2* mutant background, the protein showed both wild-type filament localization and chloroplast movements in response to whole-field blue light (Fig. 5B-C, Supplemental Movies S9 and S10). We were not able to detect any phosphopeptides associated with serine 170 with mass spectrometry analysis of THRUMIN1:YFP in transgenic *A. thaliana* in light or dark conditions (Supplemental Excel). These observations provide the first evidence we are aware of to suggest that THRUMIN1 localization to filamentous actin may not be required to achieve normal chloroplast movements.

**Figure 5.**
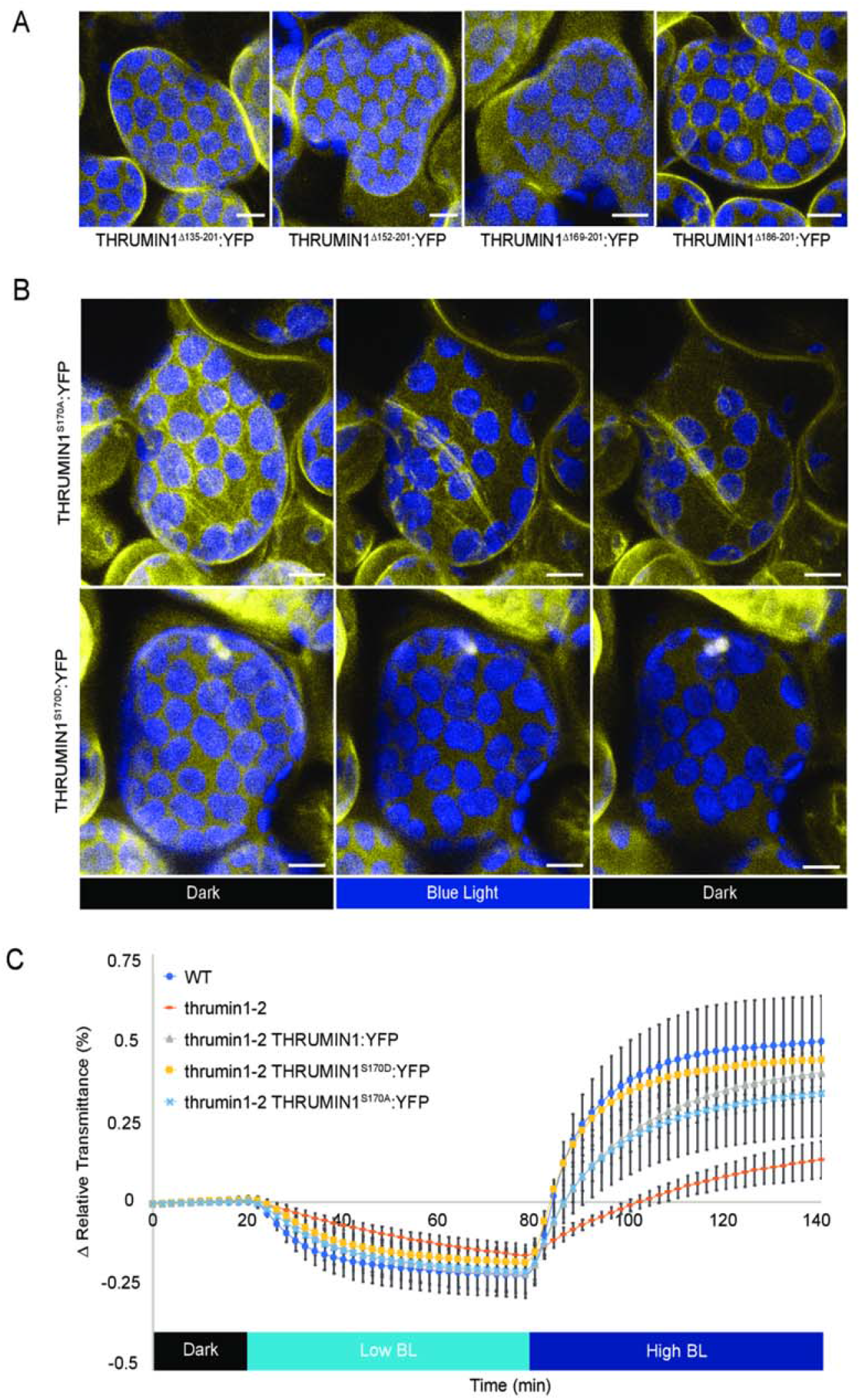
Phosphomimetic THRUMIN1 at serine 170 disrupted the filamentous localization but conferred wild type chloroplast movements. (A) Agrobacterium strains containing internal deletion constructs of the IDR were transiently expressed in *N. benthamiana* to test the localization pattern of THRUMIN1:YFP in response to blue light stimulation. Deletion of amino acids 186-201 restored the filamentous localization of THRUMIN1. (B) Serine 170, a putative phosphorylation site between amino acids 169 and 186, was mutated to alanine or aspartic acid to mimic a non-phosphorylatable or constitutively phosphorylated residue, respectively. Transient expression of 35S:THRUMIN1^S170D^:YFP in *N. benthamiana* disrupted the filamentous localization of THRUMIN1 while 35S:THRUMIN1^S170A^:YFP did not. Representative time-lapse images are shown for dark treatment (514nm for YFP excitation), blue light stimulation (470nm and 514nm), and then dark again. The scale bar indicates a 5 µm distance. (C) Leaf transmittance assays testing the response to low and high blue light of Col-0 wild type, *thrumin1-2* mutant, 35S:THRUMIN1^S170A^:YFP rescue transgenic, and the 35S:THRUMIN1^S170D^:YFP rescue transgenic. Both mutant transgenic genotypes exhibited wild type chloroplast movements. After establishment of the baseline level of leaf light transmittance after dark acclimation, chloroplast movement was induced by treatment with low blue light (∼2 µmol m^−2^s^−1^) followed by high blue light intensity (∼100 µmol m^−2^s^−1^). Standard deviation error bars represent the variance in transmittance values for 8-12 individual plants per genotype.

### Serine 113, 115, and 164 phosphorylation status do not disrupt THRUMIN1:YFP localization to F-actin

Boex-Fontvieille et al. (2014) found that THRUMIN1 was more phosphorylated in the dark and phosphorylation mutants of THRUMIN1 serine 113, 115, and 164 showed that a THRUMIN1^S113/115/164A^ triple alanine mutant failed to rescue the chloroplast movement defects in *thrumin1-2* mutant plants. The constitutively phosphorylated form of that mutant (THRUMIN1^S113/115/164D^) however, partially rescued the *thrumin1-2* mutant phenotype suggesting a reliance on a balance in phosphorylation dynamics of those sites in the light vs. dark states of THRUMIN1. When we transiently expressed YFP-tagged versions of those mutant constructs in *N. benthamiana*, neither THRUMIN1^S113/115/164A^:YFP nor THRUMIN1^S113/115/164D^:YFP disrupted THRUMIN1’s F-actin localization (Fig. 6A). Moreover, when we conducted chloroplast movement assays in *A. thaliana thrumin1-2* mutant plants expressing the THRUMIN1^S113/115/164A^:YFP transgene, we found that it rescued the chloroplast movement phenotype (Fig. 6B). Our results suggest that if phosphorylation of serine 113, 115, and 164 regulates THRUMIN1 activity, it may involve a complicated balance of phosphorylation states among those residues, possibly also involving serine 170 (Fig. 1) and/or other previously unannotated phosphorylation sites.

**Figure 6.**
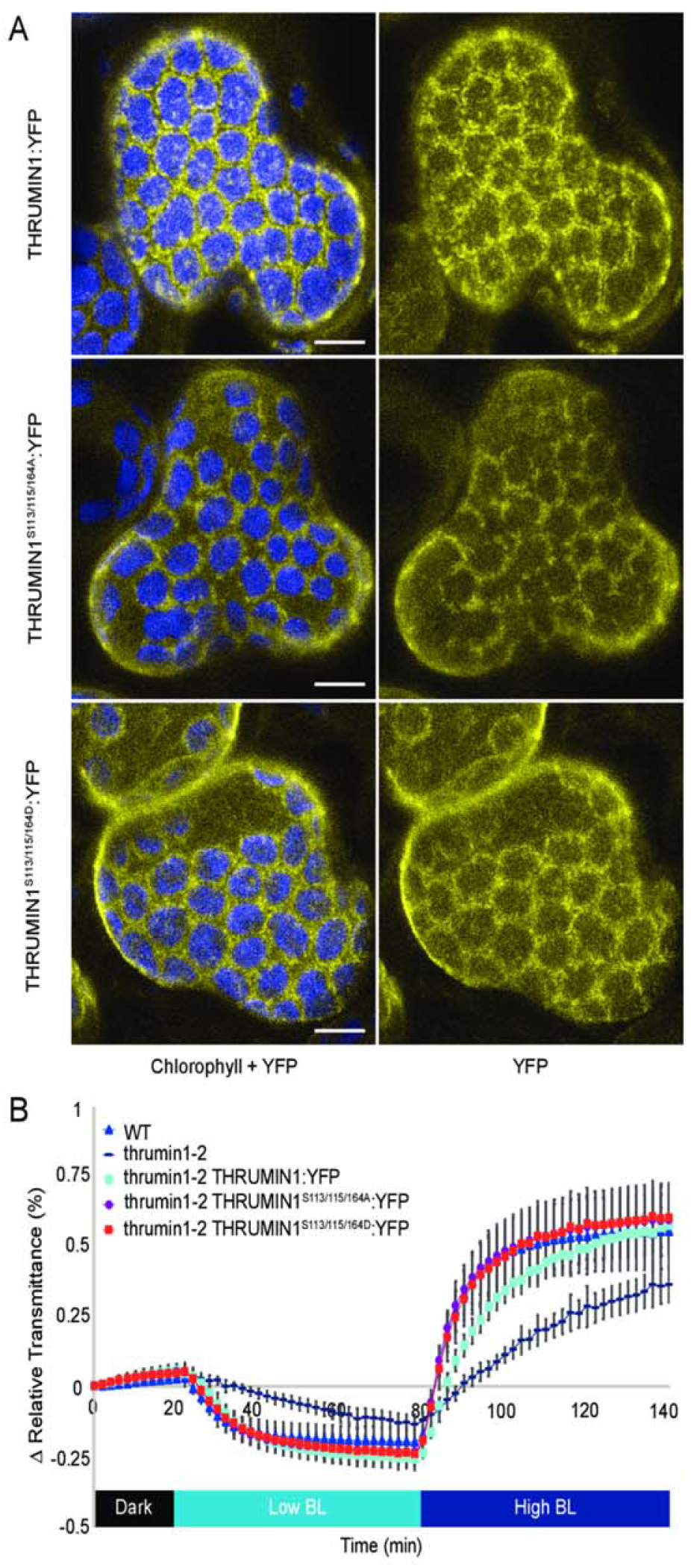
Constitutively phosphorylated and dephosphorylated THRUMIN1 at serine 113, 115, and 164 did not disrupt wild type THRUMIN1 localization or chloroplast movements. (A) Localization phenotypes of wildtype 35S:THRUMIN1:YFP and phosphorylation mutants 35S:THRUMIN1^S113/115/164A^:YFP and 35S:THRUMIN1^S113/115/164D^:YFP via transient expression in *N. benthamiana.* Both transgenic mutant plant lines phenocopied the wild type protein localization. The scale bar indicates a 5 µm distance. (B) Leaf transmittance assays demonstrated the full rescue phenotypes of 35S:THRUMIN1^S113/115/164A^:YFP and 35S:THRUMIN1^S113/115/164D^:YFP lines compared to wildtype 35S:THRUMIN1:YFP. After establishment of the baseline level of leaf light transmittance after dark acclimation, chloroplast movement was induced by treatment with low blue light (∼2 µmol m^−2^s^−1^) followed by high blue light intensity (∼100 µmol m^−2^s^−1^). Standard deviation error bars represent the variance in transmittance values for 8-12 individual plants per genotype.

### Serine 146 is required for actin filament and plasma membrane localization of THRUMIN1 in tobacco

To further examine the phosphorylation status of THRUMIN1, *A. thaliana* plants expressing the THRUMIN1:YFP transgene were treated with blue light to induce chloroplast movement and THRUMIN1 activity. Proteins were extracted and the THRUMIN1:YFP protein was immunoprecipitated with GFP-conjugated agarose beads for mass spectrometry (MS) analysis. MS analysis revealed serine 146 and threonine 104 as two new phosphorylation sites and confirmed phosphorylation at serine 113, 115, and 164. Both serine 146 and threonine 104 were mutated to phosphomimetic (aspartic acid) or non-phosphorylatable (alanine) forms. When transiently expressed in *N. benthamiana* leaf cells, mutations to threonine 104 did not alter the localization of THRUMIN1 in either the constitutively on or off forms (Supplemental Fig. S2). However, the phosphomimetic THRUMIN1^S146D^:YFP displayed cytoplasmic localization rather than wild-type filamentous protein localization, suggesting phosphorylation of serine 146 is involved in THRUMIN1 association with actin filaments as well as the plasma membrane (Fig. 7A, Supplemental Movie S11). In contrast, transient expression of THRUMIN1^S146A^:YFP localized strongly to cp-actin filaments and display biased localization away from cortical actin filaments (Fig. 7A, Supplemental Movie S12). However, the same transgenes expressed in *A. thaliana* did not confer these actin localization phenotypes (Fig. 7B). In addition, THRUMIN1^S146D^:YFP and THRUMIN1^S146A^:YFP were both able to rescue chloroplast movements in the *A. thaliana thrumin1-2* mutant background as shown by leaf transmittance assays (Fig. 7C). When the same transgenes were expressed in the Col-0 WT, wild-type localization with actin filaments was also observed indicating that the tobacco phenotype was not due to competition with the tobacco THRUMIN1 homolog (Fig. 7B). The different results observed in *N. benthamiana* and *A. thaliana* suggest that if serine 146 plays a role, it may do so in conjunction with other sites within THRUMIN1 or that it requires an unidentified factor that differs between the two species.

**Figure 7.**
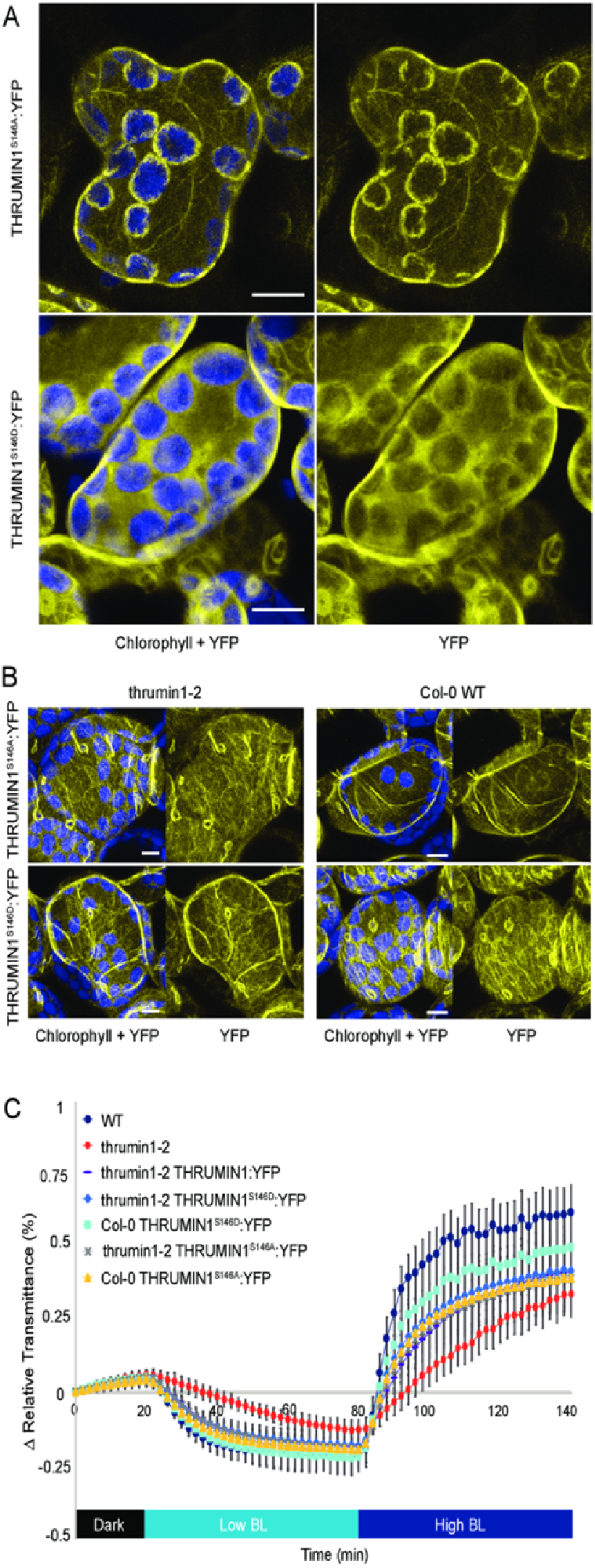
Phosphorylation mutants for THRUMIN1 displayed altered localization in *N. benthamiana* but not *A. thaliana*. (A) 35S:THRUMIN1^S146A^:YFP and 35S:THRUMIN1^S146D^:YFP transiently expressed in *N. benthamiana* showed filamentous and cytoplasmic localization, respectively. (B) However, in *A. thaliana,* 35S:THRUMIN1^S146A^:YFP and 35S:THRUMIN1^S146D^:YFP expressed in the *thrumin1-2* mutant background and Col-0 WT background displayed wild type localization patterns. The scale bar indicates a 5 µm distance. (C) Leaf transmittance assays of the mutant transgenic lines confirmed wild type chloroplast movements relative to the 35S:THRUMIN1:YFP rescue line. After establishment of the baseline level of leaf light transmittance after dark acclimation, chloroplast movement was induced by treatment with low blue light (∼2 µmol m^−2^s^−1^) followed by high blue light intensity (∼100 µmol m^−2^s^−1^). Standard deviation error bars represent the variance in transmittance values for 8-12 individual plants per genotype.

### THRUMIN1 associates with 14-3-3 proteins KAPPA and OMEGA in phosphorylation-dependent manners

Preliminary yeast-2-hybrid experiments in our lab indicated that THRUMIN1 interacted with 14-3-3 KAPPA. Our mass spectrometry experiments to assess the phosphorylation status of THRUMIN1 also led to immunoprecipitation of KAPPA with THRUMIN1. Since 14-3-3 proteins typically bind to phosphorylated residues to confer a signal (Tzivion et al., 2001), we hypothesized that KAPPA associates with one or more of the known phosphorylation sites on THRUMIN1. Co-immunoprecipitation (Co-IP) assays using non-phosphorylatable THRUMIN1 variants revealed a loss of KAPPA association when both serine 113 and 115 were mutated to alanine. Additionally, KAPPA association with THRUMIN1 was lost when the phosphomimetic mutant THRUMIN1^S113/115D^ was used as bait (Fig. 8A). Because constitutively phosphorylated or dephosphorylated states of serine 113 and 115 interfered with the ability of KAPPA to associate with THRUMIN1, the sites likely work dynamically.

**Figure 8.**
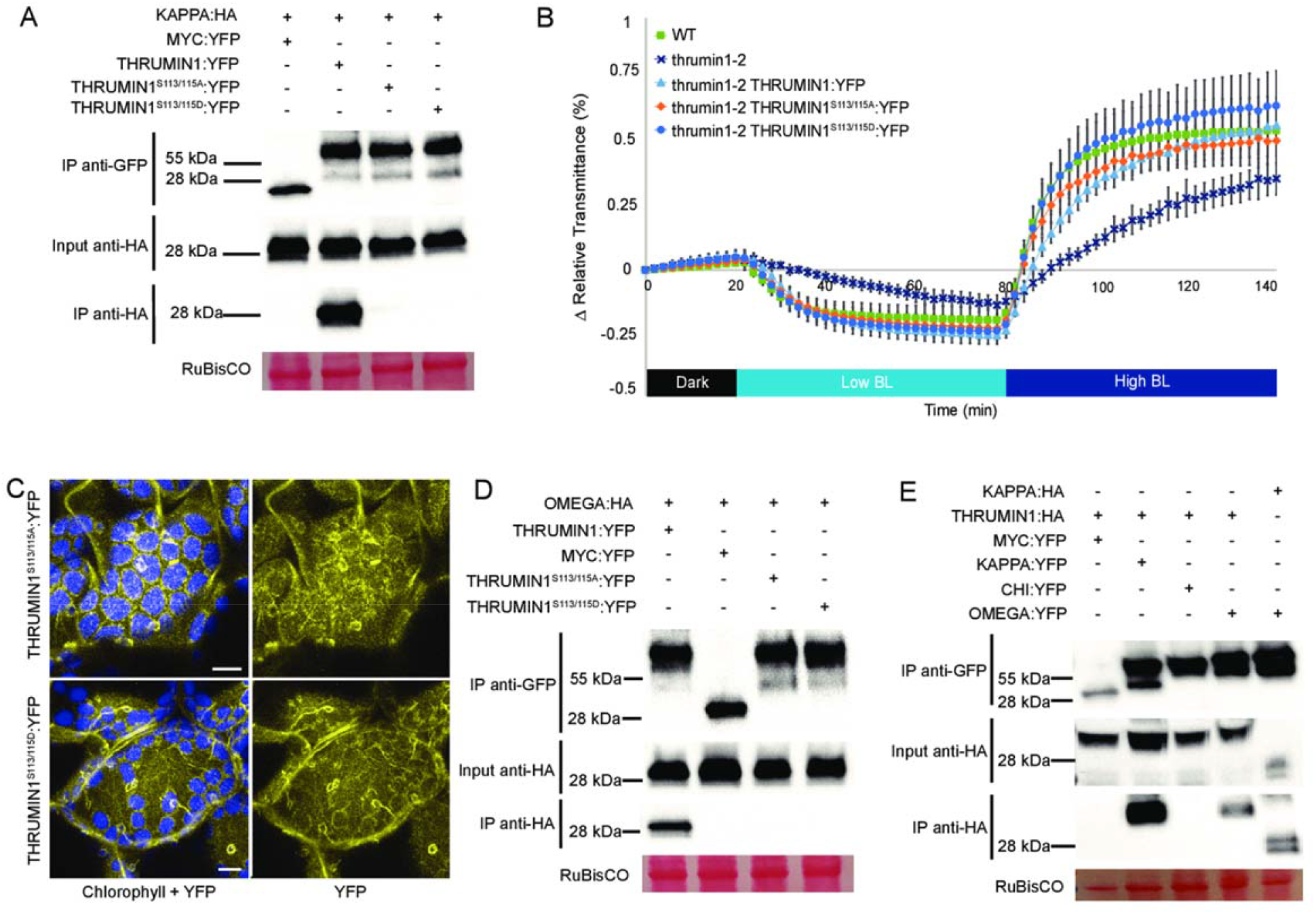
THRUMIN1 associates with 14-3-3 KAPPA and OMEGA in phosphorylation-dependent manners. (A) THRUMIN1 phosphorylation mutants THRUMIN1^S113/115A^:YFP and THRUMIN1^S113/115D^:YFP were transiently expressed under the 35S viral promoter in *N. benthamiana* and did not co-immunoprecipitate with 35S:KAPPA:HA. Wild type 35S:THRUMIN1:YFP was used as a positive control and 35S:MYC:YFP was used as a negative control. (B) Leaf transmittance assays of THRUMIN1^S113/115A^:YFP and THRUMIN1^S113/115D^:YFP transgenic plants in the *thrumin1-2* rescue background exhibited wild type chloroplast movements and (C) localization patterns. After establishment of the baseline level of leaf light transmittance after dark acclimation, chloroplast movement was induced by treatment with low blue light (∼2 µmol m^−2^s^−1^) followed by high blue light intensity (∼100 µmol m^−2^s^−1^). Standard deviation error bars represent the variance in transmittance values for 8-12 individual plants per genotype. For the micrographs, chlorophyll autofluorescence is false-colored blue and the YFP channel is separated. The scale bar indicates a 5 µm distance. All of the THRUMIN1 constructs were under control of 35S promoter. (D) Mass spectrometry analysis revealed the association of 14-3-3 OMEGA with THRUMIN1 leading to validation by *in vivo* co-immunoprecipitation in *N. benthamiana*. 14-3-3 OMEGA also bound to THRUMIN1 in a phosphorylation-dependent manner using the same serine 113 and 115 transgenic mutants as bait. (E) To verify 14-3-3 specificity, a co-immunoprecipitation assay using a different 14-3-3 protein fused with YFP as bait, 14-3-3 CHI, was tested for THRUMIN1 association in *N. benthamiana*. 35S:MYC:YFP served as a negative control while 35S:KAPPA:YFP served as a positive control. 35S:THRUMIN1:HA associates with 35S:KAPPA:YFP and 35S:OMEGA:YFP, but not 35S:CHI:YFP. 35S:OMEGA:YFP and 35S:KAPPA:HA demonstrated a typical 14-3-3 heterodimerization. All protein samples were extracted 48 hours post-infiltration. Ponceau-S stain was used as a loading control for total protein as demonstrated by RuBisCO.

When THRUMIN1^S113/115A^:YFP was expressed in *A. thaliana*, there was no obvious change in the localization of THRUMIN1 with actin filaments or chloroplast movement (Fig. 8B-C). In addition, expression of THRUMIN1^S113/115D^:YFP in the *thrumin1-2* mutant fully rescued the chloroplast movement phenotype and displayed localization with actin filaments (Fig. 8B-C). Our mass spectrometry experiments had also pulled down 14-3-3 OMEGA with THRUMIN1 and we found that the 14-3-3 OMEGA association is also dependent on the phosphorylation status of serine 113 and 115 (Fig. 8D). To test whether the THRUMIN1-KAPPA and THRUMIN1-OMEGA relationships were specific among the many 14-3-3 proteins, Co-IP assays were performed using the 14-3-3 CHI isoform. We found that THRUMIN1:HA associates with KAPPA:YFP and OMEGA:YFP, but not CHI:YFP. We also found that OMEGA:YFP and KAPPA:HA demonstrated a typical 14-3-3 heterodimerization (Fig. 8E). Although the functional relationship of these 14-3-3 proteins with THRUMIN1 remains unclear, they may play a role in light-dependent chloroplast movements by providing a scaffold by which THRUMIN1 interacts with other chloroplast movement proteins.

### Coordination of putative zinc-binding cysteines is required for light-dependent chloroplast movements and THRUMIN1-actin interactions

Previous studies in which the C-terminus of THRUMIN1 was truncated showed that it was necessary for functional chloroplast movement (Whippo et al., 2011). The C-terminus contains highly conserved clusters of cysteines with canonical zinc-binding arrangements. To determine if the coordinated cysteines are required for THRUMIN1 function, cysteines 317, 320, 351, and 354 were mutated to alanine to disrupt the putative zinc-binding capabilities. Analysis of chloroplast movements in the *thrumin1-2* mutant plants expressing THRUMIN1^C317/320/351/354A^:YFP showed that it was unable to rescue the *thrumin1-2* mutant, demonstrating the importance for these cysteine residues in chloroplast movement (Fig. 9A). However, expression of the transgene resulted in increased localization of THRUMIN1 with cp-actin filaments (Fig. 9B, Supplemental Movie S13). Upon exposure to blue light, the localization of wild-type THRUMIN1 to cp-actin typically initially disappeared but then quickly reappeared in a biased arrangement towards the leading edge of movement (Fig. 2, Supplemental Movies S1 and S2). Upon blue light exposure of dark-acclimated wild-type plants expressing THRUMIN1^C317/320/351/354A^:YFP, its localization dynamics were similar to wild-type with the exception that when the filaments reappeared, they showed a more robust and uniform arrangement around the entire periphery of the chloroplast in comparison to the wild-type protein (Fig. 9B, Supplemental Fig. S3A, Supplemental Movie S13). There was no significant change (p=0.0575) in the ratio of THRUMIN1^C317/320/351/354A^:YFP fluorescence on the leading edge versus the lagging edge during the subtle chloroplast movements (Supplemental Fig. S3B). The lack of biased relocalization and the inability of THRUMIN1^C317/320/351/354A^:YFP to rescue normal chloroplast movements adds additional support to models (Kong et al., 2013) that involve a role of biased positioning of cp-actin in regulating chloroplast movement.

**Figure 9.**
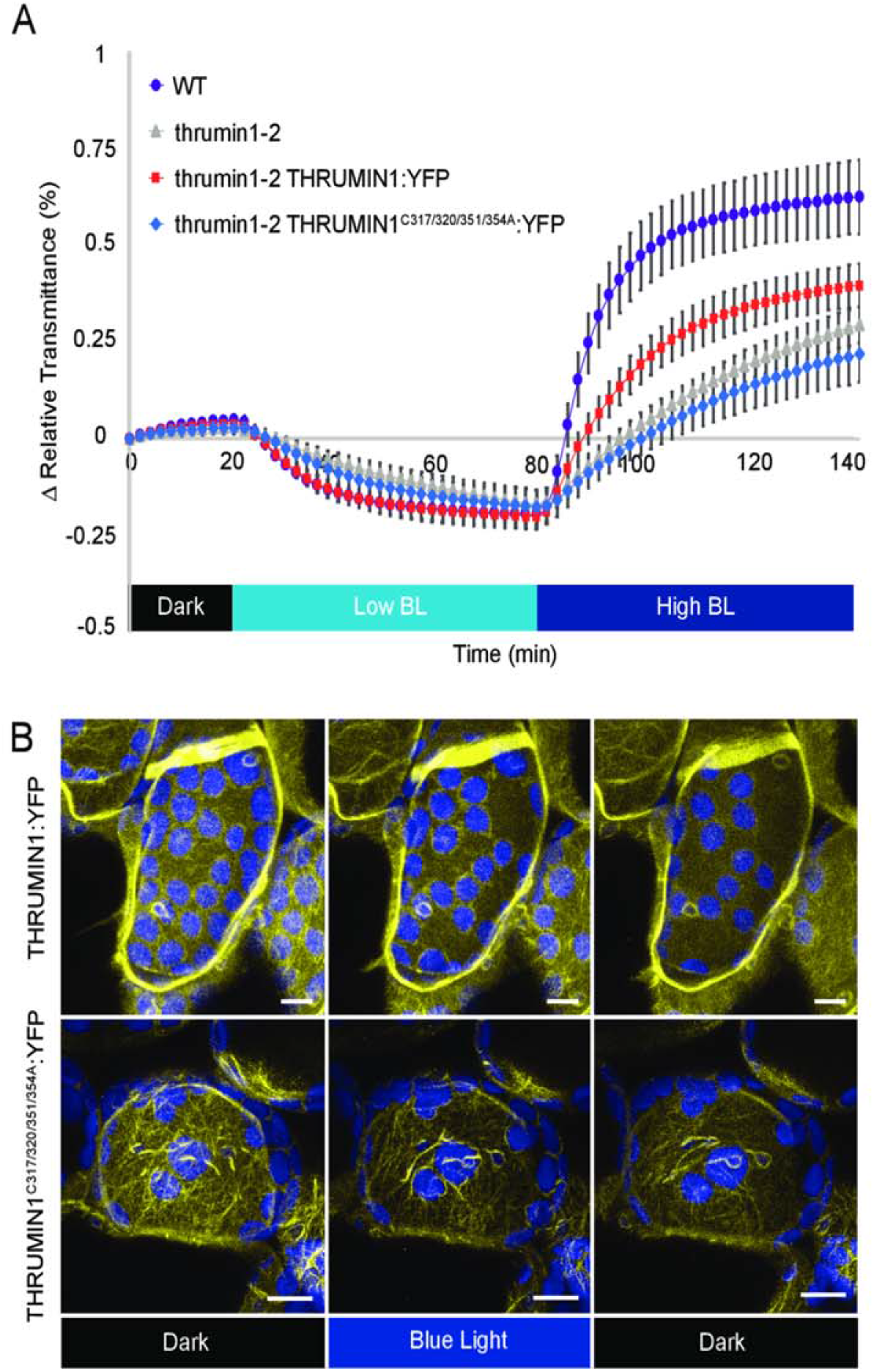
Conserved cysteine residues regulate THRUMIN1 filament reorganization and proper chloroplast movement. (A) Plants expressing 35S:THRUMIN1^C317/320/351/354A^:YFP in the *thrumin1-2* mutant background phenocopied the defective chloroplast movements assayed by leaf transmittance. After establishment of the baseline level of leaf light transmittance after dark acclimation, chloroplast movement was induced by treatment with low blue light (∼2 µmol m^−2^s^−1^) followed by high blue light intensity (∼100 µmol m^−2^s^−1^). Standard deviation error bars represent the variance in transmittance values for 8-12 individual plants per genotype. (B) 35S:THRUMIN1^C317/320/351/354A^:YFP expressed in *thrumin1-2* mutant plants displayed filamentous localization in a dominant-negative manner. In response to blue light, the filaments did not properly rearrange to facilitate effective chloroplast movement thus giving overall slower net movement. Representative time-lapse images are shown for dark treatment (514nm for YFP excitation), blue light stimulation (470nm and 514nm), and then dark again. Chlorophyll autofluorescence is false-colored blue and the YFP channel is false-colored yellow. The scale bar indicates a 5 µm distance.

## Discussion

We have conducted site-directed mutagenesis studies to obtain further insights into how the IDR associates with actin filaments in a light-dependent manner and to determine whether the THRUMIN1-actin association was directly involved in facilitating chloroplast movements. Previous work showed that the actin-binding activity of THRUMIN1 resided in the N-terminal IDR of THRUMIN1 (Whippo et al., 2011). A predicted WASP Homology-2 (WH2) domain was identified within the IDR suggesting it may be where THRUMIN1 interacts with actin filaments (Fig. 1). However, deletion of the putative WH2 domain and mutations of conserved amino acids in the WH2 domain did not disrupt the localization of THRUMIN1 to actin filaments (Fig. 4B-C). This may be because more than one WH2 domain is usually required to form actin bundles in conjunction with coordination from other actin-binding motifs (Loomis et al., 2006) or that post-translational modifications or other protein-protein associations are involved. Examination of truncations within the IDR of the full-length protein revealed a peptide region critical for the filamentous localization of THRUMIN1 (Fig. 5A) and led us to examine a putative phosphorylation site (serine 170) in that region of the protein. When serine 170 was changed to aspartic acid to mimic constitutive phosphorylation, the normal filamentous localization of THRUMIN1 was disrupted and THRUMIN1 displayed a diffuse localization pattern (Fig. 5B). However, our mass spectrometry analysis did not reveal any phosphopeptides associated with serine 170 suggesting that phosphorylation of serine 170 was too labile for our protein analysis or that the impact of this mutation may be the result of a structural change that alters the folding of the IDR interfering with THRUMIN1’s ability to bind to actin filaments. That *thrumin1-2* mutant plants expressing THRUMIN1^S170D^:YFP exhibited normal chloroplast movements (Fig. 5C) without showing localization to actin filaments was unexpected since all previous studies with THRUMIN1 and *thrumin1* mutant plants showed a close correlation between light-dependent THRUMIN1-actin interactions and chloroplast movement. These new results therefore suggest that the THRUMIN1 actin-bundling activity may not be directly involved in chloroplast movement.

Light-dependent phosphorylation of THRUMIN1 and several proteins involved in chloroplast movement (Boex-Fontvieille et al., 2014) along with the fact that the phototropins are kinases strongly suggests involvement of phosphorylation in regulating chloroplast movement. Indeed, Boex-Fontvieille et al. (2014) showed that expression of THRUMIN1 containing non-phosphorylatable alanine mutations to three phosphorylation sites that they had identified (S113, S115, S164) failed to rescue the *thrumin1-2* chloroplast movement mutant phenotype. However, when we created the same construct fused with YFP to understand how these mutations would affect the localization activity of THRUMIN1, we found no disruption to the light-induced actin-bundling characteristics (Fig. 6A) and that the mutant transgene was able to rescue the chloroplast movement defects in *thrumin1-2* mutant plants (Fig. 6B). To better assess the phosphorylation of THRUMIN1, we conducted mass spectroscopy analyses of immunoprecipitated THRUMIN1:YFP and identified phosphopeptides that contained threonine 104 and serine 146. Mutation of those amino acids to non-phosphorylatable and phosphomimetic amino acids showed that threonine 104 did not alter chloroplast movements or actin-localization with either mutated form (Supplemental Fig. S2) but mutation of serine 146 to aspartic acid completely abrogated the light-induced bundling activity of THRUMIN1:YFP (Fig. 7A). Unlike the THRUMIN1^S170D^:YFP localization, it appeared to confer a cytoplasmic localization since time-lapse confocal microscopy showed vacuolar movements in the YFP channel, which is a hallmark of cytoplasmic localization (Fig. 7A, Supplemental Movie S11). This suggests that the phosphomimetic mutation at serine 146 may in some way interfere with insertion of THRUMIN1 into the plasma membrane, possibly by affecting the N-myristoylation and S-palmitoylation groups. If so, this suggests that THRUMIN1 may be able to move between plasma membrane-localized and cytoplasmic populations. Also, because serine 146 resides close to the putative WH2 domain, it may influence the activity of the WH2 domain even though the truncations and mutations we made to the WH2 domain did not disrupt the typical THRUMIN1-actin localization. However, the results we observed in *N. benthamiana* differed from what we observed in *A. thaliana* where the transgenic plants did not confer abnormal localization of THRUMIN1 (Fig. 7B). The different results in the two species may indicate differences in how THRUMIN1 is regulated in the two model plants.

Because phosphorylation plays a critical role in the localization pattern of THRUMIN1, it is likely that proteins other than actin associate with THRUMIN1 to regulate its light-dependent function. 14-3-3 proteins are known to bind to phosphorylated targets and modulate the function of the protein. In addition, the 14-3-3 LAMBDA isoform was previously found to bind to Phototropin2 and to regulate blue light-induced stomatal opening responses via phosphorylation (Tseng et al., 2012). Although 14-3-3 LAMBDA did not have a significant effect on chloroplast movements, *A. thaliana* has 13 isoforms of 14-3-3 proteins suggesting the potential for a network of functions and/or redundancies (Rosenquist et al., 2000; DeLille et al., 2001).

The multiplicity of 14-3-3 isoforms make’s it challenging to use genetic approaches to identify individual functions of each 14-3-3 protein. In preliminary studies, we observed that 14-3-3 KAPPA interacts with THRUMIN1 via yeast-two-hybrid assays and the association was confirmed via *in vivo* co-immunoprecipitation of THRUMIN1:YFP expressed *N. benthamiana* (Fig. 8A). T-DNA insertions in the *KAPPA* locus did not cause any defects in low and high light chloroplast movements, presumably due to redundant function of other 14-3-3 proteins (Tseng et al., 2012). In this study, we identified a 14-3-3 recognition motif in THRUMIN1 that includes serine’s 113 and 115 and immunoprecipitation studies showed that changing serine 113 and 115 to either an alanine or aspartic acid disrupted the association with KAPPA and OMEGA (Fig. 8A, D) suggesting a role of serine 113 and 115 in facilitating dynamic protein-protein associations through combinatorial phosphorylation dynamics rather than a simple on or off change of a specific phosphorylation state. Our finding that 14-3-3 KAPPA and OMEGA both associated with THRUMIN1 *in vivo* suggests that they may provide scaffolding for THRUMIN1 to interact with other chloroplast movement proteins.

The cysteine-rich region in the C-terminus of THRUMIN1 was previously shown to be required for functional chloroplast movements (Whippo et al., 2011). Cysteines in that region of THRUMIN1 show canonical arrangements typical for coordination with zinc molecules (Pace and Weerapana, 2014) and suggested the cysteine clusters may be needed for proper chloroplast movement by conferring structure, facilitating protein-protein interactions, and/or playing a regulatory role. When several of the clustered cysteines were mutated to alanine’s, we saw that these sites are necessary for THRUMIN1 to undergo light-induced changes in its filamentous localization and proper chloroplast movements (Fig. 9A-B, Supplemental Fig. S3, Supplemental Movie S13). When expressed in the *A. thaliana thrumin1-2* mutant, we observed that the transgene interfered with the polar organization of filaments normally seen near the leading edge of moving chloroplasts (Fig. 9B, Supplemental Fig. S3, Supplemental Movie S13). The resulting localization of THRUMIN1 around the entire periphery instead of just near the leading edge normally seen in response to blue light suggests that controlled anchoring of chloroplasts to the plasma membrane via THRUMIN1 may be important for regulating their movement in response to blue light.

In addition to our site-directed mutagenesis work, we also used expression of the stroma-localized tpFNR:YFP probe to obtain additional insight into how the chloroplast outer envelope behaves in relation to light-dependent chloroplast movements. The contrast between the YFP signal and chloroplast autofluorescence allowed us to observe that highly dynamic, pseudopod-like protrusions of the chloroplast envelope associated with chloroplast movements (Fig. 3A,C, and Supplemental Movies S3 and S4). Consistent with Kong et al (2013), we observed that in wild-type *A. thaliana,* the chloroplast envelope protruded more in the direction that chloroplasts were moving (Supplemental Movie S3 and S4). However, *thrumin1-2* mutant plants expressing tpFNR:YFP did not develop significant protrusions of the chloroplast envelope and moved in more sporadic jumps compared to wild-type plants (Fig. 3B,C, Supplemental Movies S5 and S6). Since THRUMIN1 appears to localize along the leading edge during movement (Fig. 2, Supplemental Fig. S1, Supplemental Movies S1 and S2) our observations with tpFNR:YFP suggest that THRUMIN1 may function in bridging the chloroplast membrane to the plasma membrane and may be important for providing a footing for cp-actin dynamics to develop a motive force. The time-lapse movies of the membrane protrusions in moving chloroplasts seem to suggest a pulling mechanism but actin dynamics are generally involved in development of a pushing force. Because cp-actin is cytoplasmic and attached to the chloroplast envelope via CHLOROPLAST UNUSUAL POSITIONING1 (CHUP1) (Oikawa et al., 2003; Oikawa et al., 2008; Schmidt von Braun and Schleiff, 2008; Kadota et al., 2009), the cp-actin may anchor to the plasma membrane slightly over the chloroplast so that as the cp-actin grows away from CHUP1 on the chloroplast envelope, it can push against the plasma membrane. Although more work is needed to determine the nature of the motive force, the stroma-localized tpFNR:YFP could be a useful new tool for that work.

Current models for light-dependent chloroplast movement suggest that light activation causes cp-actin to form on the leading edge of the chloroplast and that the cp-actin in some way provides a motive force for movement via dynamic changes of the cp-actin filaments (Kadota et al., 2009; Kong et al., 2013). THRUMIN1 is a plasma membrane-localized protein that is required for normal light-induced chloroplast movements and binds to and can bundle F-actin (Whippo et al., 2011). Moreover, wild-type THRUMIN1:YFP has been found to localize with cp-actin in leaf mesophyll cells in a light-dependent manner and has led us and others to suggest it plays a role in motive force dynamics (Whippo et al., 2011; Kong et al., 2013). In this study we found that mutating serine 146 and serine 170 to aspartic acid disrupted THRUMIN1-actin filament localization but did not seem to interfere with chloroplast movement dynamics (Figs. 5, 7 and Supplemental Movies S7, S8, S11) suggesting that THRUMIN1’s interaction with cp-actin may not be directly associated with the motility mechanism as previously thought. Furthermore, mutations in the C-terminal cysteine residues resulted in the loss of THRUMIN1’s normal polar localization at the leading edge of cp-actin filaments after blue light exposure and instead relocalized to cp-actin around the entire chloroplast periphery to disrupt chloroplast movements (Fig. 9, Supplemental Fig. S3, Supplemental Movie S13). Also, in the *thrumin1-2* mutant, chloroplasts exhibit sporadic jumps of movement without any discernable chloroplast protrusions of the chloroplast envelope (Supplemental Movies S5 and S6) instead of the more sustained movements and envelope protrusions seen in wild-type leaf cells (Supplemental Movies S3 and S4). Taken together, these findings lead us to propose that THRUMIN1 functions more to anchor chloroplasts to the plasma membrane rather than regulating the motive force for chloroplast movements. The initial detachment of THRUMIN1 from cp-actin may thus function to release the chloroplasts so that they can move via cytoplasmic streaming, or another motive force. Upon moving, reorganization of the THRUMIN1-actin association along the leading edge may then serve to reattach the chloroplast to the plasma membrane and guide the path of movement through its interactions between cp-actin and the plasma membrane.

## Materials and Methods

### Growth Conditions

All seeds were surface sterilized with 19:1 87.5% ethanol 30% H_2_O_2_. After drying, the seeds were plated in Petri dishes on ½ MS salts and 0.6% agar and cold stratified for 3-days at 4°C. The stratified seeds were then grown for 10 days under ∼100 µmol m^−2^s^−1^ white light in growth chambers with 12-hour photoperiods at 23°C then transplanted to fertilized (20-20-20) potting mix (PromixB) and grown under ∼120 µmol m^−2^s^−1^ white light and a 12-hour photoperiod. The same process was used for transgenic lines except the ½ MS media contained 30 mg/mL BASTA (glufosinate ammonium; Cayman Chemical) for selection of the transgenic plants.

### Site-directed mutagenesis and deletion constructs

Site-directed mutations of the phosphorylation sites were created using similar methods as the Quickchange® II XL kit from Agilent technologies by using the primers listed below to recreate the whole vector via PCR with the new mutation in place. The primers with either aspartic acid or alanine substitutions were used to amplify the mutated version of THRUMIN1 using the Gateway vector pBSDONR P1P4 backbone (Qi et al., 2012). Error-free sequences were recombined into the pEG100 plant expression vector (Earley et al., 2006) with a pBSDONR P4rP2 YFP clone using LR Clonase II (Invitrogen, Carlsbad, CA, USA) to create a final destination vector to be transformed into *Agrobacterium* strain GV3101. Internal deletion constructs were created using primers that extend away from the deletion site towards the beginning and end of the gene to create two PCR fragments which were later fused together through PCR extension (Atanassov et al., 2009). The fused products were recombined into pEG100 in the same manner as described earlier. The mutant genes were then transformed into *A. thaliana* Col-0 and *thrumin1-2* (SALK_027277) backgrounds using the *Agrobacterium-*mediated floral dip transformation method (Clough and Bent, 1998).

### *Agrobacterium*-mediated Transient Expression

*Agrobacterium* (strain GV3101) carrying the different gene constructs in the pEG100 plant expression vector were cultured in LB media and resuspended in 10mM MgCl_2_ 10mM MES pH 5.6 to an OD_600_ of 0.2. The solution was incubated for several hours with 3′,5′-Dimethoxy-4′-hydroxyacetophenone (Acetosyringone) to induce virulence and then injected into *N. benthamiana* leaves. After 48 hours of incubation, leaves were excised and mounted for imaging the fluorescence of the expressed gene products on a Leica SP8 scanning confocal microscope using imaging parameters as described below.

### Live Cell Confocal Microscopy

Prior to mounting leaf samples on slides, the plants were low-light acclimated for ∼3 hours under ∼10 µmol m^−2^s^−1^ light intensity to facilitate arrangement of the chloroplasts on the periclinal cell face before imaging. After low-light acclimation, small leaf sections excised and mounted for imaging and incubated in Perfluoroperhydrophenanthrene (CAS Number 306-91-2, Millipore/Sigma) to clear out the air spaces and optimize image resolution. All imaging was acquired using a Leica SP8 Scanning Confocal microscope with an inverted 40x/1.10 water objective lens. During the first 5 minutes, time-lapse images of YFP fluorescence (525-600 nm) in palisade mesophyll cells were captured at continuous intervals of ∼25 seconds with only YFP excitation using 514nm laser illumination to prevent activation of the phototropin photoreceptors. The samples were then exposed for ∼15 minutes with whole-field or microbeam blue light stimulation (470nm) to induce the avoidance response while imaging YFP excitation (514nm). The blue light treatment was then stopped and the cells were imaged for an additional ∼5-10 minutes with 514nm YFP excitation in the absence of blue light stimulation. Throughout the imaging process, chlorophyll emission was also collected at 650-720nm. In all cases, the top 12 µm of the palisade mesophyll cell was imaged in 0.42 µm Z-steps. The images were combined by Z-projection and analyzed using FIJI software.

### THRUMIN1 Cp-actin Localization Quantification

To calculate THRUMIN1:YFP fluorescence intensities on the leading edge of the chloroplast versus the lagging edge, kymographs were generated in Fiji from time-lapse confocal microscopy movies using a 10 pixel-wide segmented line in the path of chloroplast movement using the KymographBuilder plugin (created by Hadrien Mary). THRUMIN1 fluorescence gray values for the leading and lagging edges were then extracted by tracing a 10 pixel-wide segmented line down the Y-axis of the kymograph. Distance values (microns) on the kymograph were converted to time from the known frame rate intervals of the time-lapse movies. The fluorescence values of the leading edge were then ratioed over the lagging edge. The ratios were averaged from at least 13 chloroplast replicates and were calculated at two minutes before the peak difference, at the peak, and 2 minutes after the peak. The statistical significance of the difference between the leading and lagging ratios before and after peak chloroplast movement was assayed by paired two-tailed T-test.

### Stromal Marker Protrusion Quantification

Chloroplast membrane protrusion events were identified by the difference between the fluorescence of the stromal maker, tpFNR:YFP, and the chlorophyll autofluorescence from confocal time-lapse movies. After a high blue light microbeam irradiation, the positions of the first chloroplast membrane protrusion at the onset of chloroplast movement were binned for the leading and lagging edges. Data were obtained from at least 30 different chloroplasts from 8 different movies.

### Mass Spectrometry and Co-Immunoprecipitation

To identify phosphorylation sites, ∼3.0 grams of transgenic *A. thaliana* plants expressing THRUMIN1:YFP were exposed to high-intensity blue light for 10 minutes and flash frozen in liquid nitrogen. The powdered plant material was mixed with lysis buffer containing 50 mM Tris-HCl pH 7.5, 150 mM NaCl, 10% Glycerol, 1 mM EDTA, 1% NP40, and plant protease inhibitor cocktail tablets (Millipore/Sigma; cOmplete™, Mini, EDTA-free Protease Inhibitor Cocktail; 11836170001) and mixed by rotation using a tube rotator at 4°C for 30 minutes. The plant lysate was then centrifuged at 10,000xg and the supernatant added to washed GFP-Trap agarose beads (Chromotek; gta20) and mixed via tube rotator at 4°C for 3 hours. After incubation with the lysate, the beads were washed five times with the lysis buffer at 4°C with 1000xg centrifugation to pellet the beads between washes. The beads were washed with a pre-urea wash buffer (100 mM ammonium bicarbonate (ABC), pH 8.0) by resuspending and rotating for 3-5 minutes at 4°C. After the wash, the beads were pelleted at 1000xg and the supernatant was removed. For protein elution, the beads were resuspended in urea buffer (8M Urea, 100mM ABC pH 8.0) at a 1:1 ratio to the bead volume and incubated for 10-30 min at room temperature with occasional mild vortex pulsing, pelleted, and the supernatant (containing eluted protein) saved in a tube. Elution off the beads was repeated two more times and the combined eluates were submitted for mass spectrometry analysis (see methods) at the Indiana University Laboratory for Biological Mass Spectrometry facility.

Co-Immunoprecipitation assays were performed similarly to the mass spectrometry protocol. Samples were extracted from ∼0.5 gram of *N. benthamiana* leaves that were transiently expressing the transgene of interest. Instead of a urea elution, the final resuspension of the washed beads was in a 1X SDS-loading buffer and boiled at 95°C for 10 minutes. Samples were centrifuged at 1000xg to pellet the beads and the supernatant was loaded into a Mini-PROTEAN TGX 4-20% (w/v) gradient gel (Bio-Rad) to run for 45 minutes at 150V. After SDS-PAGE, the protein was transferred to a nitrocellulose membrane (GE Lifescience Product #10600003) and stained with Ponceau-S for validation of protein transfer. The blots were then blocked in 5% skim milk for at least one hour. For GFP/YFP detection, primary anti-mouse GFP antibody (Novus Biological; NB600-597) was applied at 1:7500 dilution and incubated overnight at 4°C with horizontal platform shaking (62 RPM) followed by secondary goat anti-mouse-HRP antibody (A-10668; Invitrogen) incubation with three 5-minute Tris-Buffered Saline 0.1% Tween 20 (TBST) washes in-between and after antibody applications. For HA detection, anti-HA-HRP (3F10; Sigma) conjugated antibody was applied at 1:7500 dilution for 1 hour at room temperature. Blots were incubated with Clarity Western ECL Substrate (#1705061; Bio-Rad) for 5 minutes and imaged using a ChemiDoc Imaging System to detect chemiluminescence of the blotted proteins.

For mass spectrometry analysis, tryptic peptides were injected into an Easy-nLC 100 HPLC system coupled to an Orbitrap Fusion Lumosmass spectrometer (Thermo Fisher Scientific). Specifically, peptide samples were loaded onto an Acclaim PepMap 100 C18 trap column (75 μm x 2 cm, 3 μm bead size with 100 Å pores) in 0.1% (v/v) formic acid. The peptides were separated using an Acclaim PepMap RSLC C18 analytical column (75 μm x 25 cm, 2 μm bead size with 100 Å pores) using an acetonitrile-based gradient (solvent A: 0% [v/v] acetonitrile and 0.1% [v/v] formic acid; solvent B: 80% [v/v] acetonitrile and 0.1% [v/v] formic acid) at a flow rate of 300 nL/min. A 30-min gradient was performed as follows: 0 to 0.5 min, 2 to 8% B; 0.5 to 24 min, 8 to 40% B; 24 to 26 min, 40 to 100% B; 26 to 30 min, 100% B, followed by re-equilibration to 2% B. Electrospray ionization was then performed with a nanoESI source at a 275°C capillary temperature and 1.9-kV spray voltage. The mass spectrometer was operated in data-dependent acquisition mode with mass range 400 to 2000 m/z. Precursor ions were selected for tandem mass spectrometry analysis in the Orbitrap with 3-sec cycle time using higher energy collisional dissociation at 28% collision energy. The intensity threshold was set at 5 3 104. The dynamic exclusion was set with a repeat count of 1 and exclusion duration of 30 sec. The resulting data were searched in Protein Prospector (http://prospector.ucsf.edu/ prospector/mshome.htm) against the *Arabidopsis* database. Carbamidomethylation of Cys residues was set as a fixed modification. Protein N-terminal acetylation, oxidation of Met, protein N-terminal Met loss, pyroglutamine formation, phosphorylation on STY were set as variable modifications. In total, three variable modifications were allowed. Trypsin digestion specificity with one missed cleavage was allowed. The mass tolerance for precursor and fragment ions was set to 10 ppm for both. Peptide and protein identification cutoff scores were set to 15 and 22, respectively. All phosphopeptide spectra were confirmed manually to verify peptide identity and site of modification.

### Transmittance Curve Assay

Leaf discs (7mm) were made with a hole punch and placed on a 0.5% agar pad in wells of clear-bottom 96-well plate (Falcon) sealed with Microseal ‘A’ film (Bio-Rad). The film was punctured over each well with a needle to allow for gas exchange. The prepared plates were dark-acclimated for a minimum of 6 hours before placement in a BioTek Cytation 3 Imaging Reader. The baseline level of light transmittance through the leaf discs was calculated from measurements of absorbance values of 660 nm red light taken every 2 min for 20 min (red light does not activate chloroplast movement). To induce chloroplast movement in the cells, the plate reader was programmed to eject the plate for exposure to the selected light intensity for two minutes. The plate was then moved back into the plate reader for recording of transmittance values (660 nm red light absorbance) for each well. After each reading, the plate was re-ejected to return to the blue light treatment. The cycle of recording transmittance values and incubating with blue light was repeated for the indicated time periods for a given light treatment. The calculated changes in light transmittance values were normalized to be relative to the starting ‘dark’ position values.

### Primer List

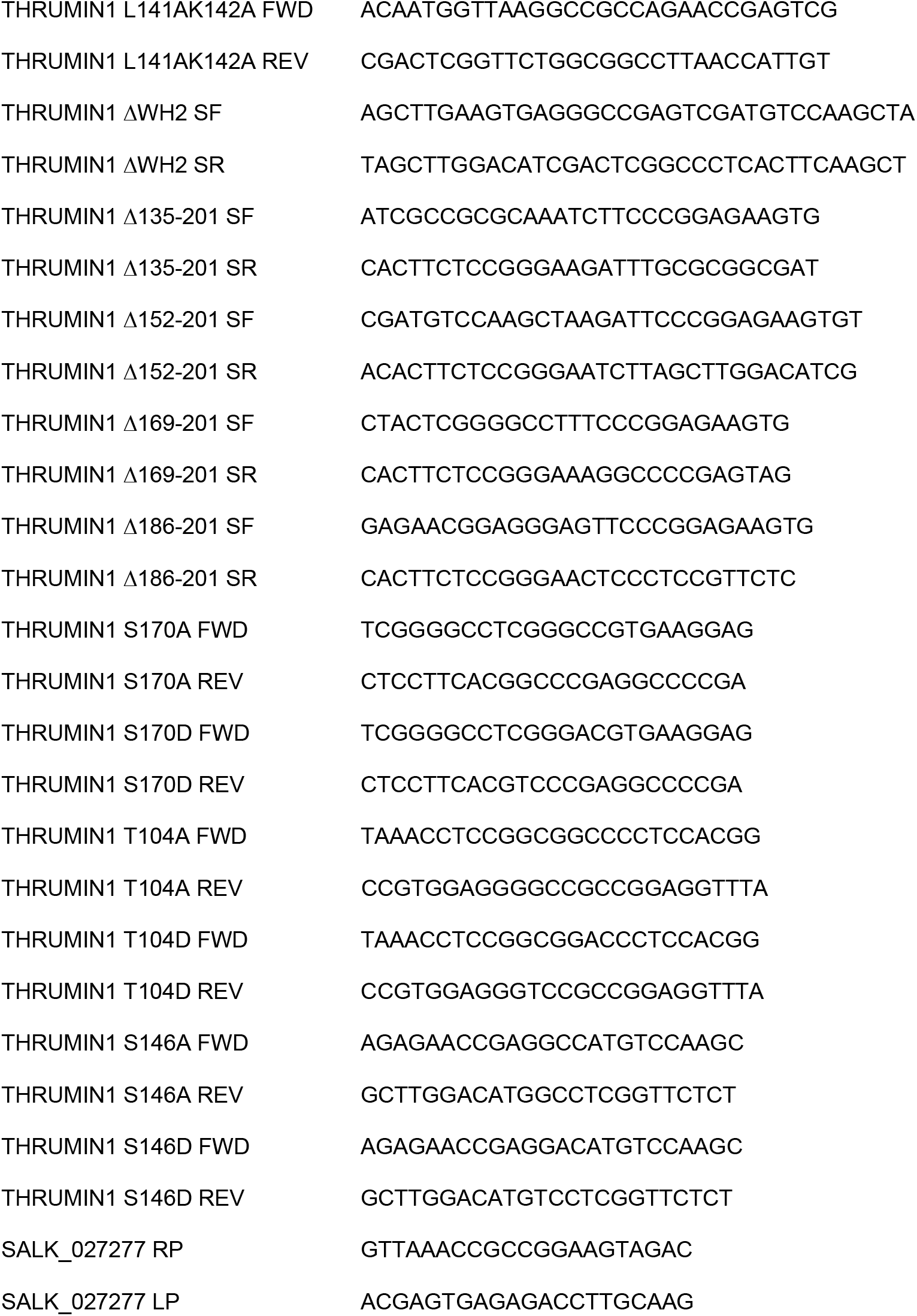
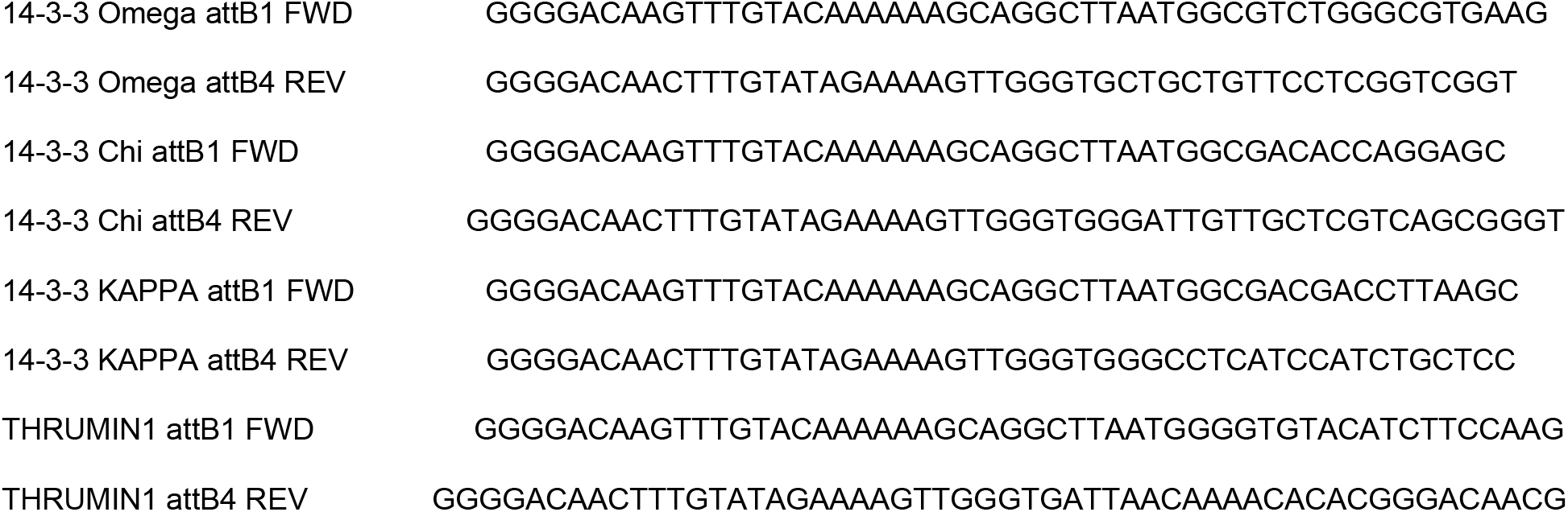

## Supplemental Material

**Supplemental Movie S1**

Irradiation of a mesophyll cell with a microbeam of 470 nm blue light (rectangle) resulted in the initial loss of 35S:THRUMIN1:YFP (yellow channel) from cp-actin around the chloroplast periphery (chlorophyll autofluorescence in blue channel) and 35S:THRUMIN1:YFP reassociation towards the leading edge as the chloroplasts exit the irradiated area.

**Supplemental Movie S2**

Exposure of a mesophyll cell to whole-field 470 nm blue light increases 35S:THRUMIN1:YFP (yellow channel) association with the periphery of chloroplasts (chlorophyll autofluorescence in blue channel) with a bias to the leading edge of those chloroplasts that moved towards the cell edge.

**Supplemental Movie S3**

Stroma-localized tpFNR:YFP (yellow channel) and chlorophyll autofluorescence (blue channel) revealed pseudopod-like membrane protrusions associated with the leading edge of chloroplast movement in response to microbeam irradiation with 470 nm blue light (rectangle). The all-yellow organelles are plastids in the overlying epidermal cells.

**Supplemental Movie S4**

Stroma-localized tpFNR:YFP (yellow channel) and chlorophyll autofluorescence (blue channel) revealed pseudopod-like membrane revealed more randomly localized pseudopod-like protrusions, and chloroplast movements, when mesophyll cells were exposed to whole-field 470 nm blue light. The all-yellow organelles are plastids in the overlying epidermal cells.

**Supplemental Movie S5**

Stroma-localized tpFNR:YFP (yellow channel) and chlorophyll autofluorescence (blue channel) in mesophyll cell of the *thrumin1-2* mutant revealed minor development of pseudopod-like membrane protrusions in response to microbeam irradiation with 470 nm blue light (rectangle). The all-yellow organelles are plastids in the overlying epidermal cells.

**Supplemental Movie S6**

Stroma-localized tpFNR:YFP (yellow channel) and chlorophyll autofluorescence (blue channel) in mesophyll cell of the *thrumin1-2* mutant revealed minor development of pseudopod-like membrane protrusions and erratic chloroplast movements in response to whole-field irradiation with 470 nm blue light (rectangle). The all-yellow organelles are plastids in the overlying epidermal cells.

**Supplemental Movie S7**

Transiently expressed 35S:THRUMIN1^S170D^:YFP in *N. benthamiana* failed to localize to filaments and conferred a more diffuse localization phenotype with or without whole-field 470 nm blue light.

**Supplemental Movie S8**

THRUMIN1^S170D^:YFP mutant in *A. thaliana thrumin1-2* mutant plants showed diffuse localization but rescued the defective chloroplast movement phenotype.

**Supplemental Movie S9**

Non-phosphorylatable THRUMIN1^S170A^:YFP mutant expressed in *A. thaliana thrumin1-2* mutant plants showed both wild-type filament localization and chloroplast movements in response to whole-field 470 nm blue light.

**Supplemental Movie S10**

Non-phosphorylatable THRUMIN1^S170A^:YFP mutant expressed in *N. benthamiana* showed both wild-type filament localization and chloroplast movements in response to whole-field 470 nm blue light.

**Supplemental Movie S11**

Transiently expressed THRUMIN1^S146D^:YFP in *N. benthamiana* showed cytoplasmic localization rather than wild-type filamentous protein localization with or without whole-field 470 nm blue light.

**Supplemental Movie S12**

Transiently expressed THRUMIN1^S146A^:YFP in *N. benthamiana* localized strongly to cp-actin filaments and display biased localization away from cortical actin filaments with or without whole-field 470 nm blue light.

**Supplemental Movie S13**

Expression of 35S:THRUMIN1^C317/320/351/354A^:YFP in the *A. thaliana thrumin1-2* mutant resulted in increased localization of THRUMIN1 with cp-actin filaments around the chloroplast perimeter and failed to rescue chloroplast movements in response to whole-field 470 nm blue light.

## Acknowledgements

We would like to acknowledge the Indiana University Light Microscopy Imaging Center for training and use of the scanning confocal microscope (Leica SP8), the Indiana University Laboratory for Biological Mass Spectrometry facility for sample analyses, and Dr. Sid Shaw for generously sharing his microscopy and image analysis expertise.

---

**Supplemental Figure S1**

THRUMIN1-cp-actin localization increased at the leading edge of *A. thaliana* palisade mesophyll chloroplasts in response to blue light. (A) Representative time-course plot of fluorescence intensities of THRUMIN1:YFP at the leading edge of a chloroplast versus the lagging edge of the chloroplast measured with KymographBuilder (see methods). Upon exposure to a high blue light microbeam (at 4 minutes), the fluorescence intensity of both the leading and lagging edge initially decreased, which was followed by an increase in THRUMIN1:YFP fluorescence intensity at the leading edge while the fluorescence intensity continued to decrease along the lagging edge. As the time course progressed, the leading edge fluorescence intensities of THRUMIN1:YFP decrease as the chloroplast moved out of the blue light microbeam. (B) Ratios of leading to lagging edge fluorescence intensities 2 minutes before the fluorescence at the leading edge reached its peak, at its peak, and 2 minutes after the peak. The data were collected from 13 chloroplasts from 7 cells. The results show that the leading/lagging-edge fluorescence ratio is statistically significant (p=0.0291). Error bars = standard deviation.

**Supplemental Figure S2**

Threonine 104 does not alter the filamentous localization of THRUMIN1. (A) *N. benthamiana* cells transiently expressing 35S:THRUMIN1^T104A^:YFP displayed no alterations to the localization pattern of wild type THRUMIN1. (B) Similarly, the phosphomimetic variant 35S:THRUMIN1^T104D^:YFP did not alter the localization of THRUMIN1 when expressed transiently in *N. benthamiana*. Representative time-lapse images are shown (514nm for YFP excitation) and chlorophyll autofluorescence is false-colored blue while the YFP channel is false-colored yellow. The scale bar indicates a 5 µm distance.

**Supplemental Figure S3**

Mutations to THRUMIN1’s conserved cysteines altered reorganization of the cp-actin at the chloroplast periphery. (A) Representative time-course plot of fluorescence intensities of leading and lagging edges of mesophyll chloroplasts in *Arabidopsis* expressing 35S:THRUMIN1^C317/320/351/354A^:YFP as measured with KymographBuilder (see methods). Upon exposure to a high blue light microbeam (at 2.6 minutes), the fluorescence intensity of both the leading and lagging edge decreased followed by an increase in both the leading and lagging edge fluorescence intensity. As the time course progressed, the leading and lagging edge fluorescence intensities of THRUMIN1^C317/320/351/354A^:YFP fluctuated and there was very little chloroplast movement as seen in Fig. 9. (B) Ratios of leading to lagging edge fluorescence intensities 2 minutes before the fluorescence at the leading edge reached its peak, at its peak, and 2 minutes after the peak. The data were collected from 15 chloroplasts from 7 cells. The results show that the leading/lagging-edge fluorescence ratio was not statistically significant (p=.0575) with the 35S:THRUMIN1^C317/320/351/354A^:YFP mutant. Error bars = standard deviation.

## Notes

**Funding Information:** This work was supported by the National Science Foundation Grant MCB-0848083 and Indiana University.

